# Calcium-dependent facilitation of P/Q-type calcium channels governs the polarity and magnitude of short-term synaptic plasticity

**DOI:** 10.64898/2026.04.24.720678

**Authors:** Shuwen Chang, Maria Gurma, Yi-Mei Yang, Lu-Yang Wang

**Author notes:** Correspondence: Dr. Yi-Mei Yang < > or Dr. Lu-Yang Wang < >. Lead Contact: Dr. Lu-Yang Wang < >.

## Abstract

P/Q-type calcium channel (Cav2.1) is the major channel that mediates Ca^2+^ influx during action potentials (APs) and evokes neurotransmitter release from presynaptic terminals. Repetitive activity induces its Ca^2+^-dependent facilitation (CDF) via binding of calmodulin (CaM) superfamily proteins to the IQ-like motif, specifically isoleucine (I) and methionine (M) sites, on the cytoplasmic c-terminus of Cav2.1. However, whether and how CDF contributes to short-term synaptic plasticity remains elusive. By recordings from the calyx of Held terminal in IQ-like motif point mutation knock-in mice (Cav2.1 IM-AA KI), we found that activity-dependent CDF is completely abolished, resulting in lower quantal output and shorter release time course as well as profound reductions in the magnitute of short-term facilitation and depression (STF and STD) in different Ca^2+^ concentrations. Prolonging deactivation of Ca^2+^ channels by broadening spike width normalizes quantal output and release time course in Cav2.1 IM-AA synapses, but does not fully rescue STF/STD. These results indicate that CDF of Cav2.1 channels governs the polarity and magnitude of short-term synaptic plasticity in fast-spiking central synapses.

## Introduction

Short-term synaptic plasticity is manifested as temporal facilitation and/or depression of synaptic strength during repetitive presynaptic activity, namely short-term facilitation (STF) and depression (STD), which influence the information transfer between neurons¹. It occurs in the time scale of milliseconds to seconds that provides an instant response to the previous stimuli and helps shape the synaptic strength during sustained activity. While different synapses display distinct polarity in the direction of short-term plasticity, some synapses exhibit both synaptic STF and STD. Over the years, multiple mechanisms have been proposed to explain STF/STD. However, the key determinants for these two forms of plasticity remain ambiguous and controversial, but are inevitably related to the dynamics of Ca^2+^ signaling in the nerve terminal.

At presynaptic terminals, Ca^2+^ influx through voltage-gated calcium channels (VGCCs) plays important roles in triggering neurotransmitter release and regulating short-term plasticity. During repetitive stimulation, the accumulation of intracellular Ca^2+^ transiently increases the release probability (Pr) of synaptic vesicles (SVs) ^1,2^. As a result, synaptic strength temporally increases, leading to STF. On the other hand, in synapses with high initial Pr, Ca^2+^ rise can lead to a rapid depletion of the readily-releasable pool (RRP) of SVs, resulting in a use-dependent decline in quantal output or STD^1^. Several hypotheses have been put forward to explain how Ca^2+^ buildup shapes synaptic strength and plasticity. These models include (1) residual Ca^2+^ from the preceding action potential (AP) preoccupies Ca^2+^ binding sites of the low affinity Ca^2+^ sensor that triggers release, thereby enhances the synaptic strength by Ca^2+^ influx from subsequent APs^3^; (2) There is a separate high-affinity Ca^2+^ sensor that independently mediates synaptic facilitation, i.e. Synaptotagmin 7^4^; (3) an endogenous Ca^2+^ buffer saturation leads to an accumulation of Ca^2+^ that increases the Pr of SVs^5^; (4) activity-dependent modification of VGCCs by incoming Ca^2+^ directly controls Ca^2+^influx into the terminal and leads to changes the synaptic strength^6^. While there are multiple studies in support of these competing ideas, unequivocal evidence in establishing the role of Ca^2+^-dependent modification of VGCCs in STF and STD during repetitive activity remains sparse.

P/Q-type VGCCs (Cav2.1) represent the major calcium channel type that mediates AP-evoked neurotransmitter release at presynaptic terminals. Unlike N-type VGCCs (CaV2.2) that often coexist in the same terminals, P/Q-type VGCCs undergo dual feedback regulation:calcium-dependent facilitation and inactivation (CDF and CDI) through their interactions with calcium-binding proteins such as calmodulin (CaM) or other CaM superfamily proteins ^2,7,8^ . This bipartite regulation is mediated by its IQ-like CaM binding domain of the intracellular C-terminus ^8,9^, where the binding of the amino acids isoleucine (I) and methionine (M) in IQ-like motif to CaM induces conformational changes that facilitate the channel opening within milliseconds^10^.

As neurotransmitter release steeply correlates to presynaptic Ca^2+^ entry, activity-dependent CDF is predicted to have profound effects on the magnitude of STF and STD. Indeed, regulation of presynaptic Ca^2+^ channels has been reported to contribute to short-term synaptic plasticity at the calyx of Held^11–14^ and cerebellar cerebellar synapses ^15^ in the central nervous system. After genetic deletion of Cav2.1 channels at the calyx of Held synapse, CDF was greatly diminished, so was pair-pulse facilitation of synaptic output ^16,17^, suggesting presynaptic CDF as the major drive for synaptic facilitation. However, complete knock out of Cav2.1 led to a compensatory expression of other VGCCs, making it difficult to study the independent effect of P/Q type Ca^2+^ channel facilitation on synaptic plasticity. To address the question of how CDF shapes synaptic plasticity, Cav2.1 IM-AA knock-in (KI) mice, in which native Cav2.1 are replaced by mutant Cav2.1 with paired alanine (A) substitutions for isoleucine (I) and methionine (M) residues in the IQ-like motif (IM-AA), have been generated ^18–20^.

Using the IM-AA KI mice, Nanou et al^6^ showed that STF is largely attenuated at the neuromuscular junction, hippocampal CA3-CA1 synapse and CA3-basket cell synapse in the IM-AA mice. In contrast, Weyrer et al^21^ showed that STF at CA3-CA1 synapse was not affected and even increased in cerebellar parallel fiber (PF)-Purkinje cell (PC) and PC-deep cerebellar nucleus (DCN) synapses. However, the lack of direct analyses of VGCCs at the presynaptic terminals renders it unintelligible to resolve the issues of whether and how IM-AA mutation regulates Ca^2+^ channel properties at the nerve terminals by CaM superfamily proteins and causally influences STF and STD in native synapses.

To overcome the limitation, we use the calyx of Held synapse in the auditory brainstem, where Cav2.1 channels contribute to 99% of presynaptic Ca^2+^ current after the postnatal 2 weeks ^22,23^. Owing to its large size, the calyx of Held synapse offers an optimal experimental model to directly study the amplitude, voltage-dependent gating kinetics, CDF and CDI of presynaptic P/Q calcium channels and their impact on glutamate release and short-term plasticity. We found that the IQ-like motif is not only important for activity-dependent CDF but also crucial for the channel targeting and gating at the presynaptic terminal. As a result, IM-AA KI synapse displayed a significantly reduced Ca^2+^ current density, shortened transmitter release time course of SVs evoked by a single AP. Furthermore, IM-AA mutation completely abolishes CDF, and dramatically alters the patterns and magnitude of STD and STF in different extracellular calcium concentrations. Our results demonstrate that the activity-dependent CDF actively shapes the patterns of short-term synaptic plasticity, which is potentially important for expanding the dynamic range and fidelity of neurotransmission, particularly during intense neural activity.

## Results

### IQ motif regulates the density of P/Q-type VGCCs in presynaptic nerve terminals

Taking advantage of direct access to presynaptic terminals of mature calyx of Held in which P/Q-type calcium channels exclusively mediate glutamate release, we measured the currents mediated by VGCCs (I_Ca_) in postnatal (P) 16 days to P21 littermate WT and IM-AA KI mice. The I_Ca_ were elicited by 20 ms voltage steps ranging from -100 mV to +50 mV (Figure 1 A, B, D, E). To quantitatively compare the density of calcium channels per terminal, we normalized the peak I_Ca_ to the membrane capacitance and displayed as I_Ca_ density. The relationships between peak or tail I_Ca_ density are illustrated as a function of voltage in Figure 1C, F. Unexpectedly, the I_Ca_ density C was reduced by 35% compared with WT (Figure 1C; −87.76 ± 7.85 pA/pF vs. −59.17 ± 6.01 pA/pF; WT: n = 16, IM-AA KI: n = 8; t test, *p = 0.0259). I_Ca_ current density was fitted with a Boltzmann function, revealing no changes in half-activation or kinetics in IM-AA KI synapses (Figure 1F; WT: V_1/2_ = -23.53 mV, k = 6.68; IM-AA KI: V_1/2_ = -23.48 mV, k = 7.93). These results suggest that the abundance of functional P/Q-type VGCCs is reduced at calyces of IM-AA KI mice.

**Figure 1:**
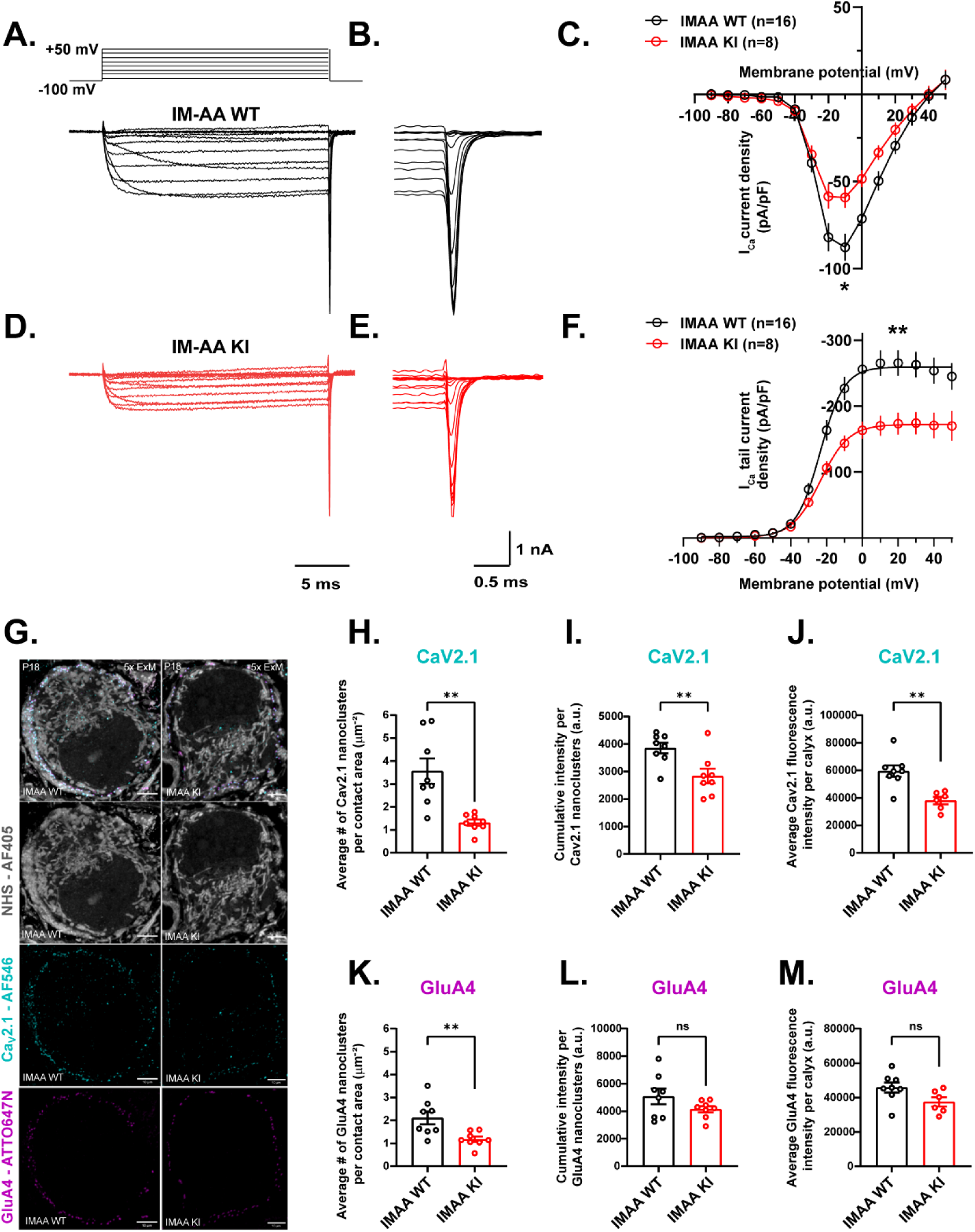
IQ motif regulates calcium channel density and gating in presynaptic nerve terminals。. **(A, B, D, E)** Representative traces presynaptic Ca^2+^ current (ICa) evoked by 20-ms depolarizing voltage steps from a holding potential of -90 mV to test potentials +50 mV (in 10mV steps) in calyceal synapses from P16-P21 WT (black) and IM-AA KI (red). **(C)** steady-state and **(F)** peak tail current-voltage (I-V) relationship of Ca2+ currents density for WT (black circles) and IM-AA KI (red circles) synapses. Dashed lines indicate Boltzmann fits with V1/2 of - 23 mV and slope factor k=6.7 for WT and V1/2 for -23 mV with k=7.9 for IM-AA KI. (**G)** Representative ∼5x expansion microscopy (ExM) images of calyces from P18 WT and IM-AA KI mice labeled with NHS (AF405, grey), GluA4 (ATTO647N, magenta) or Cav2.1 a1-subunits (AF546, cyan) antibodies. NHS-ester (grey) pan-protein stain shows ultrastructural context of the calyx to distinguish presynaptic and postsynaptic compartments. Scale bars: 10 µm (**H**) Scatter plots of the average Cav2.1 nanocluster number per contact area (WT: 3.562 ± 0.5673 μm^−2^, IM-AA KI: 1.315 ± 0.057 μm^−2^) (**I**) Scatter plots of the cumulative fluorescence intensity per cluster (correlate of Cav2.1 number per cluster), expressed as absolute value (WT: 3851 ± 331, IM-AA KI, 2835 ± 331). (**J**) Scatter plot of average Cav2.1 fluorescence intensity in a single calyx (WT: 59381 ± 5393, IMA-AA KI: 38050 ± 5393). (**K**) Scatter plots of the average GluA4 nanocluster number per contact area (WT: 2.110 ± 0.3029 μm−2, IM-AA KI: 1.178 ± 0.3029 μm−2). (**L**) Scatter plots of the cumulative fluorescence intensity per cluster (correlate of GluA4 number per cluster), expressed as absolute value (WT: 5074 ± 611.2, IM-AA KI: 4144 ± 611.2). (**M**) Scatter plot of average GluA4 fluorescence intensity in a single calyx (WT: 45803 ± 4148, IMA-AA KI: 37495 ± 4148). Data are shown as the means ± SEMs. Statistical significance was determined by t test. **p<0.01; ns, not significant.

To directly visualize subsynaptic distribution patterns of VGCCs with high resolution, we next applied ∼5x expansion microscopy (ExM)^24^ to co-label P/Q-VGCCs (Cav2.1-AF546, cyan) and GluA4 AMPAR subunits (GluA4-ATTO647N, magenta) in P18 IM-AA WT and KI synapses (2-3 calyces each mouse, 3 pairs of WT and IM-AA mice). The GluA4 labelling was used to confirm Cav2.1 nanoclusters at active zones, as each active zone is associated with juxtaposed AMPA receptor (AMPAR) nanocluster on the postsynaptic density, also known as a nanocolumn^25^. Pan-protein staining of the hydrogel-embedded tissues (NHS-AF405, grey) provided ultrastructural context to help examine the protein distributions in the calyx pre- and postsynaptic compartments. 3D nanocluster analysis of each calyx enabled quantitative comparison of Cav2.1 and GluA4 puncta between two genotypes as depicted in Figure 1G. Under the same threshold conditions for two sets of images, the average Cav2.1 nanocluster density and total fluorescence intensity in IM-AA calyces was substantially fainter and more diffusive than that in WT calyces (Figure 1I, J), being reduced by ∼30% and ∼36% respectively, in line with functional measurements of I_Ca_ current density (Fig 1A-F). The GluA4 nanocluster density and overall fluorescence intensity was not significantly different between the two genotypes (Fig 1L, M), indicating that presynaptic nanodomain integrity is disrupted in IMAA calyces while postsynaptic scaffolding is preserved. Interestingly, the average total number of both Cav2.1 and GluA4 nanoclusters was reduced in IM-AA calyces (Fig 1H, K), suggesting a reduced number of functional synaptic release sites and nanocolumns. These data implicate the important role of IQ motif in regulating the targeting and/or trafficking Cav2.1 channels to presynaptic terminals as well as Ca^2+^ entry to impact transmitter release.

### Reduction in presynaptic I_Ca_ in IM-AA KI mice attenuates the amplitude and decay time course of evoked EPSCs

Because the amount of neurotransmitter released is steeply dependent on presynaptic Ca^2+^concentrations^1^, we further tested the functional impact of attenuated I_Ca_ on synaptic strength in IM-AA KI synapses. By stimulating afferents with a bipolar electrode, we recorded AP-evoked excitatory postsynaptic currents (eEPSCs) from the principal neurons in the medial nucleus of the trapezoid body (MNTB) of P16-21 WT and IM-AA KI mice. Indeed, the amplitude of eEPSCs from KI synapses was reduced by 32% compared to that from the WT (Figure 2A-B, WT: -10.83 ±0.63 nA, n=17; KI: -6.89 ± 0.58 nA, n=23; t-test, **** p<0.0001). Surprisingly, we found that the half-width of eEPSCs of IM-AA KI synapses were reduced (Figure 2E, WT : 0.67 ± 0.02 ms, n=17 versus KI: 0.62 ± 0.01 ms n=23; K-S test, * p=0.01) accompanied by an accelerated decay time constant (Figure 2D, WT: 0.51 ± 0.02 ms, n=17 versus KI: 0.47 ± 0.01 ms, n=23; t-test, * p=0.04) and slightly increased rise time (Figure 2C, WT : 0.24 ± 0.01 ms, n=17 versus KI: 0.26 ± 0.01 ms, n=23;K-S test, ns p=0.23). We thus analyzed the miniature EPSCs (mEPSC) to rule in or out the possible contributions from changes in the kinetics or number of postsynaptic AMPA receptors. Both mEPSC amplitude (Figure 2F-G, WT: 53.87 ± 3.19 n=8 versus KI: 54.61 ± 2.31 pA n=9; t-test, ns, p=0.85) and frequency (Figure 2J, WT: 3.40 ± 0.58 n=8 versus KI: 5.79 ± 1.23 n=9; t-test, ns, p=0.1122) were not significantly different between IM-AA WT and KI. However, the rise (Figure 2H, rise time WT : 0.14 ± 0.004 ms n=8 versus KI: 0.16 ± 0.004 ms n=9; t-test,*p=0.0218) and decay time constants (Figure 2I, WT: 0.24 ± 0.08 ms n=7 versus KI: 0.28 ± 0.08 ms n=9; t-test,**p=0.0045) of the mEPSC were slightly but significantly slower in the IM-AA KI mice compared to the WT. These data suggest slower rise time of eEPSCs may be attributed to AMPAR kinetics, in line with the observation of ExM results showing no significant reduction in cumulative intensity of GluA4 nanoclusters in IM-AA calyces (Figure 1G-M). The shorter half-width of eEPSC in IM-AA KI synapses cannot be accounted for by postsynaptic mechanisms. If anything, the slower mEPSC kinetics might partially offset the faster SV release rate in IM-AA KI mice. These data suggest the faster time course of eEPSCs in IM-AA synapses must be due to presynaptic mechanisms, most likely by shortened Ca^2+^ transients when the density and nanocluster number of P/Q-type channels was reduced in IM-AA calyces (Figure 1).

**Figure 2:**
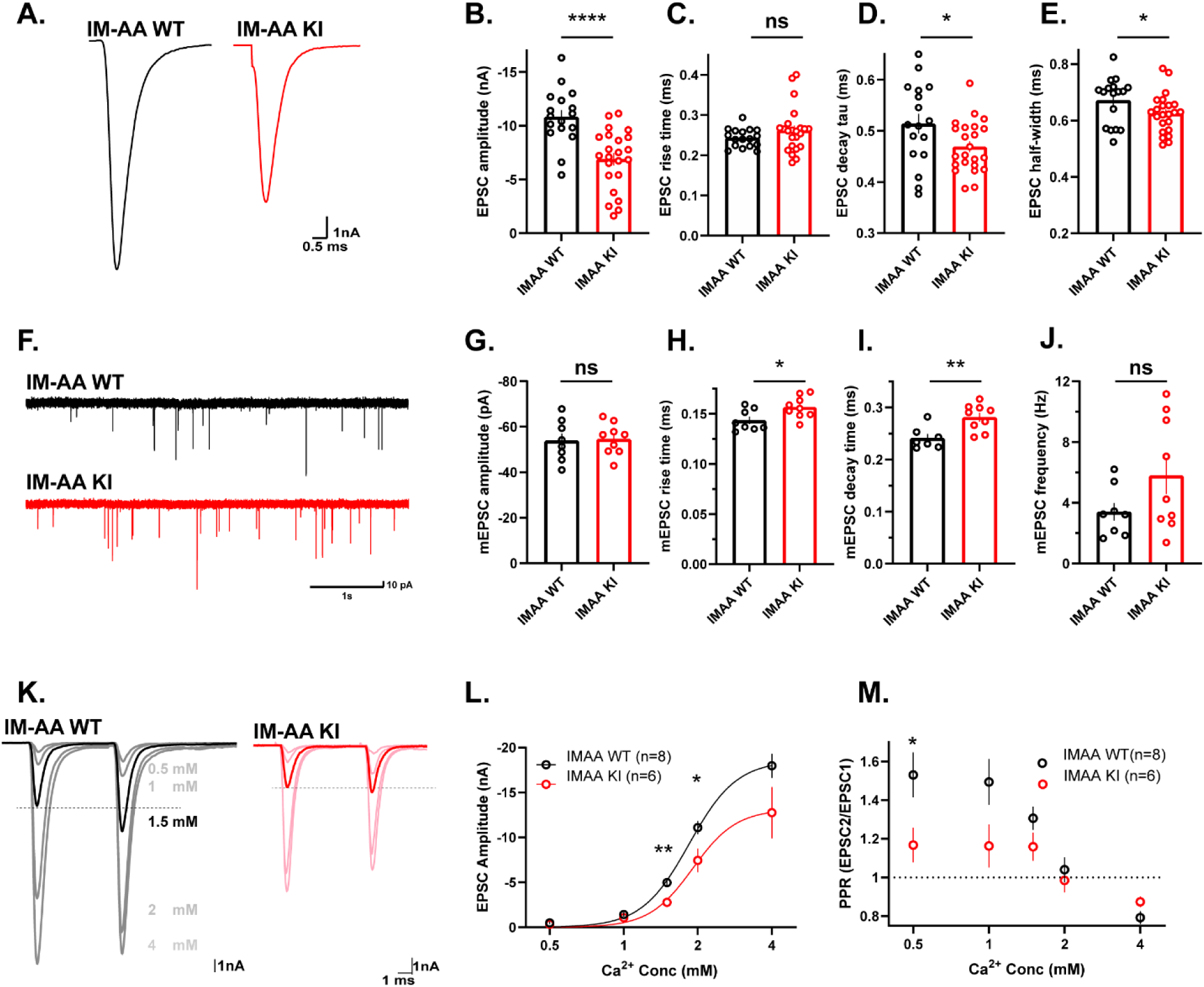
Reduced presynaptic I_Ca_ in IM-AA KI mice decreases evoked EPSC amplitude and accelerates EPSC decay without altering quantal properties. Evoked EPSCs were elicited by afferent fiber stimulation and recorded under voltage-clamped principal neurons of the MNTB regionof P16-P21 WT and IM-AA KI mice. **(A)** Representative EPSC traces from WT (black) and IM-AA KI (red) recorded in 2 mM external Ca^2+^. **(B-E)** Scatter plots summarizing eEPSC **(B)** amplitude, **(C)** rise time, **(D)** decay time, and **(E)** half-width for WT (black) and IM-AA KI (red) mice. **(F)** Representative mEPSC traces from a WT (black) and an IM-AA KI (red) synapse. **(G-J)** Scatter plots of the mEPSC **(G)** amplitude, **(H)** rise time, **(I)** decay time, and **(J)** frequency. **(K)** Representative traces of eEPSCs recorded from WT (black) and IM-AA KI (red) synapses at external Ca^2+^ concentrations of 0.5, 1, 1.5, 2 and 4 mM.(L) Ca^2+^ concentration-response relationship of the eEPSC amplitudes for WT (black) and IM-AA KI (red) synapses. **(M)** Ca^2+^ dependence of paired-pulse ratio (PPR) for WT (black) and IM-AA KI (red) synapses. Data are shown as the means±SEMs. Statistical significance was determined by t test or K-S test. ****P<0.0001; **p<0.01; *p<0.05; ns, not significant.

To further explore variables in the presynaptic locus, we subsequently studied the effect of altering the driving force for Ca^2+^ influx on the synaptic strength in WT and IM-AA KI synapses by probing the amplitude (Figure. 2K-L) and pair-pulsed ratio (PPR; Figure. 2M) of eEPSCs at different external Ca^2+^ concentrations ([Ca^2+^]_o_: 0.5, 1, 1.5, 2 and 4 mM Ca^2+^). The amplitudes and PPR were plotted as a function of [Ca^2+^]_o_ in a logarithmic scale. We found that the power relation between [Ca^2+^]_o_ and release is not significantly different between WT (h=4.5, n=8, R=0.92) and IM-AA KI synapses (h=4.9, n=6, R=0.68). However, the eEPSC amplitudes were consistently smaller in IM-AA KI at different [Ca^2+^]_o_ (1 mM: WT:-1.43 ± 0.22 versus KI: -1.07 ± 0.19 n=8 and 6 respectively, t-test, **p=0.0057 ; 2 mM: WT: -11.09 ± 0.72 versus KI: -7.45 ± 1.28, t-test, *p=0.0209 ; 4 mM Ca^2+^ WT: -17.98 ± 1.32 versus KI: -12.75 ± 2.84 nA, t-test, ns p=0.09), with no change in apparent EC50, indicating the sensitivity of release sensor to Ca^2+^ was not affected. However, we noted the higher [Ca^2+^]_o_, the larger the difference in the amplitude of eEPSCs. Conversely, the lower [Ca^2+^]_o_, the bigger the PPR difference between two genotypes. In other words, pair-pulse depression could be converted to pair-pulse facilitation as [Ca^2+^]_o_ lowers in WT, but this increase was less prominent in IM-AA KI synapses. These consistent changes of eEPSC amplitudes and PPRs in IM-AA KI synapses implicate the density of Ca^2+^ channels as the key variable being altered by IM-AA mutations, giving rise to the phenotypical differences in presynaptic Ca^2+^ influx, quantal output and short-term plasticity.

### IM-AA mutation attenuates activity- and Ca^2+^-dependent facilitation of presynaptic I_Ca_

Mutations at two residues (IM) in IQ motif of Cav2.1 have previously been shown to attenuate CDF and enhance CDI in recombinant cell-line expression systems and in soma of neurons^6,9^. However, it is not known whether and how the IQ motif affects use-dependent properties of I_Ca_ mediated by P/Q-type channels in central nerve terminals. To directly address this question at the calyx of Held synapse, we further compare the changes of presynaptic I_Ca_ in WT and IM-AA KI synapses in response to pseudo-AP trains. I_Ca_ was evoked with 200 ms pulse trains mimicking AP waveform (0.3 ms from V= -80 mV to +40 mV) at frequencies of 100 and 300 Hz. The I_Ca_ peak charge were measured and plotted as a function of time and normalized to the first I_Ca_ charge. As shown in Figure 3A, E, I_Ca_ facilitated in a highly frequency-dependent manner in the WT synapse without use-dependent inactivation. The magnitude of I_Ca_ facilitation reached the highest level of ∼125% at 300 Hz within the shortest time (Figure 3D, WT 300 Hz: 124 ± 6 % n=11). Similar but less pronounced changes were observed at 100 Hz (Figure 3C, 100 Hz: 112 ± 2% n=15). To further test the Ca^2+^ dependency of CDF, we measured I_Ca_ in the 1 mM external Ca^2+^. Interestingly, we found the degree of CDF at 1 mM Ca^2+^ is comparable to 2 mM Ca^2+^ concentration (Figure 3F, WT 300 Hz:121±7% n=5). Conversely, IM-AA mutants’ synapses exhibited no I_Ca_ facilitation at any given frequencies (Figure 3B, D, E; KI 300 Hz: 102 ± 2 % n=12; and Figure 3C, 100 Hz: 104 ± 1% n=13). Given that CDF is mediated by Ca^2+^-dependent association of CaM superfamily proteins with P/Q-type VGCCs, our results demonstrate that the IQ motif of Cav2.1 is necessary for CDF of I_Ca_ in central terminals.

**Figure 3:**
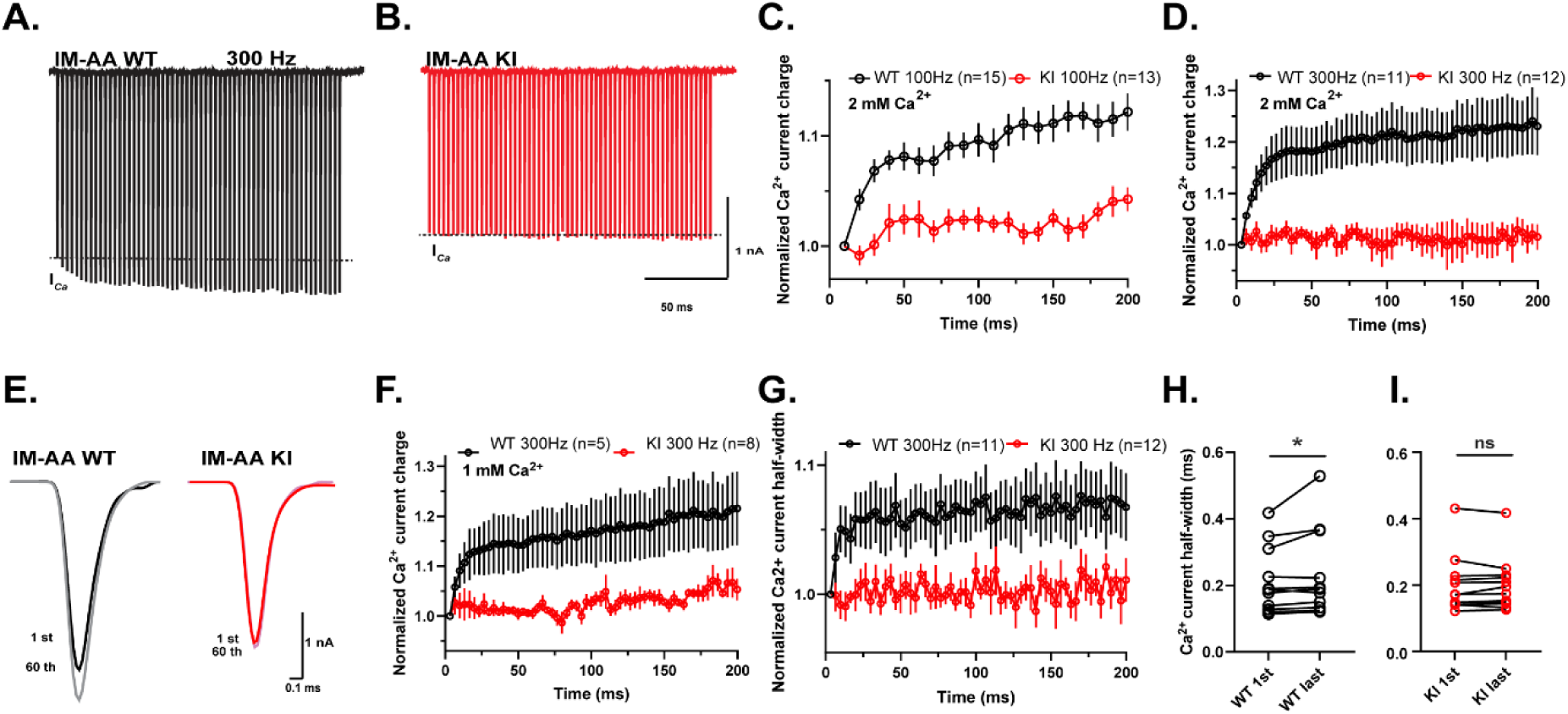
IM-AA mutation attenuates activity- and Ca^2+^-dependent facilitation of presynaptic I_Ca_. **(A, B)** Representative presynaptic Ca^2+^ current (I_Ca_) recorded during 200 ms trains of presynaptic depolarizations delivered at 300 Hz (0.3-ms steps from V_h_=-80 mV to + 40 mV) in WT (black) and IM-AA KI (red) calyces. **(C, D)** Pooled data of I_Ca_ integral plotted as a function of time at 100 Hz and 300 Hz respectively from WT (black) and IM-AA KI (red). **(E)** Superimposed traces of the first I_Ca_ and last (33 rd) ICa responses from the 300 Hz train shown at an expanded time scale. **(F, G)** Pooled data of I_Ca_ charge **(F)**and half-width **(G)** plotted as function of time during the stimulus train. **(H, I)** Scatter plot comparing the half-width of the 33^rd^ (last) I_Ca_ during 300 Hz train in **(H)** WT and **(I)** IM-AA KI calyces. Data are shown as the means ± SEMs. Statistical significance was determined by Wilcoxon matched pairs rank test or paired t-test. *p<0.05; ns, not significant.

Because I_Ca_ facilitation is highly frequency dependent, we postulated that there might be activity-and Ca^2+^-dependent changes in the gating of P/Q-type Ca^2+^ channels. We thus analyzed the kinetic changes of I_Ca_ during the trains. Interestingly, we found I_Ca_ from WT synapses displayed a significantly longer decay time constant at the first pseudo-AP and further prolonged during the 300 Hz train (Figure 3G, H, 107 ± 3% n=11, *p=0.02), suggesting there might be pre-association of CaM superfamily proteins with IQ-like motif, giving rise to a difference in basal deactivation of I_Ca_. On the other hand, there were no changes in IM-AA mutants at any given frequencies (Figure 3G, I, 300 Hz:101 ± 2% n= 12, ns, p=0.88). These data raise the possibility that there might be an activity-dependent recruitment and binding of CaM superfamily proteins to the IQ-motif and/or Ca^2+^ dependent configuration change, similar to the model previously proposed for CaM N- and C-lobe switching in L-type VGCCs^26^, to allow full manifestation of CDF during sustained neuronal activities.

### IM-AA mutation changes the magnitude of short-term facilitation and depression

At any given synapse, the number of SVs released during a single AP is a product of Pr and the size of RRP of SVs. To estimate the RRP, we measured the cumulative amplitudes of eEPSCs during a 200 ms, 300 Hz train in both genotypes (Figure 4A-B), as STD at the calyx of Held is largely caused by depletion of the RRP^27,28^. In Figure 4C, peak amplitudes of eEPSC during 300Hz train are summed to give a plot of cumulative eEPSC amplitudes. The last 5-10 data points (steady state) were fitted by linear regression and back-extrapolation to time zero. The zero time intersect gives an estimate of the size of RRP. We found the RRP size between WT and IM-AA KI mice was not significantly different (Figure 4D; WT: -49.54±5.14 nA n=10 and IM-AA KI: -47.56±8.44 nA n=13, p=0.85, t-test, ns, p=0.8545). To estimate Pr, we divided the amplitude of the first eEPSC by the estimated RRP in WT and IM-AA KI synapses. As expected, IM-AA KI synapses showed lower Pr compared to WT synapses (Figure 4E; WT: 19.79±3.4 % n=10 versus KI: 13.81±1.2 % n=13, t-test, ns, p=0.086).

**Figure 4:**
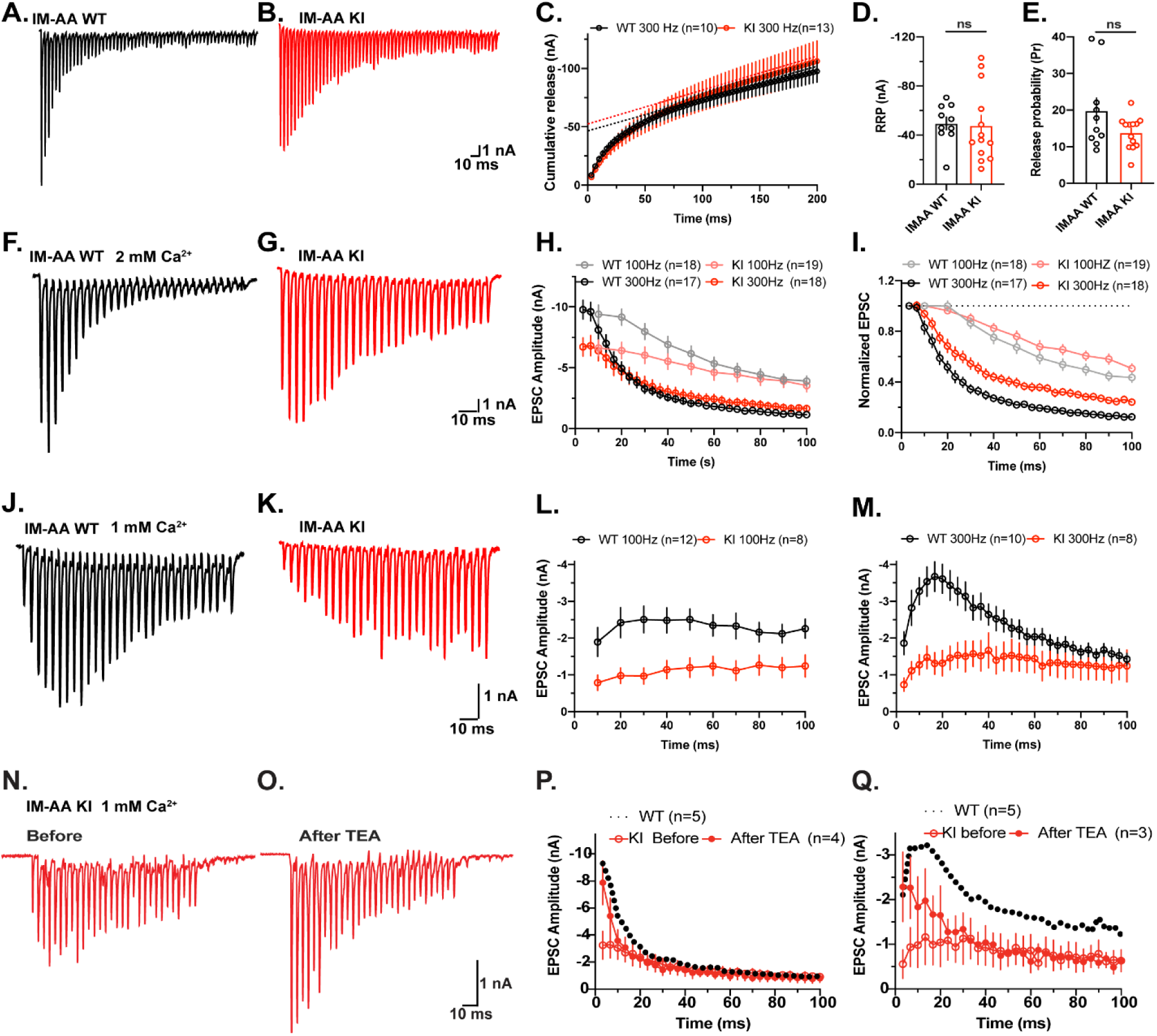
IM-AA mutation alters the magnitude of short-term facilitation and depression, which is normalized by slowing I_Ca_ deactivation with TEA. **(A, B)** Representative eEPSCs of WT (black) and IM-AA KI (red) synapsesduring 200 ms trains of APs delivered at 300 Hz in 2 mM external Ca^2+^. **(C)** Cumulative eEPSC amplitudes during 300 Hz stimulation for WT (black) and IM-AA KI (red) mice. The last 10 points were fitted by linear regression and extrapolated to time zero to estimate the readily releasable pool (RRP) size. **(D, E)** Scatter plots showing the estimated RRP size **(D)** and (**E)** vesicular release probability of the synaptic vesicle (Pr) for WT (black) and IM-AA KI (red) synapses. **(F, G)** Representative eEPSCs from WT (black) and IM-AA KI (red) synapses elicited by 100-ms trains of 300Hz AP trains at 2 mM external Ca^2+^ and **(J, K)** 1 mM external Ca^2+^. **(H, L)** Pooled eEPSC amplitudes and **(I, M)** normalized eEPSC amplitudes plotted as a function of time during 100, 300Hz for WT (black) and IM-AA KI (red) synapses. eEPSCs amplitudes were normalized to the first eEPSC in each train. **(N, O)** Representative eEPSCs from IM-AA KI synapses elicited by 100-ms trains of 300 Hz APs before **(N)** and after **(O)** application of 50 μM TEA-Cl at 1 mM external Ca^2+^. **(P)** eEPSC amplitudes plotted as a function of time at 300Hz for IM-AA KI synapses before and after TEA-Cl application in 2 mM Ca^2+^; the dashed line represents WT responses in 2 mM external Ca^2+^. **(Q)** eEPSC amplitudes plotted as a function of time in 1 mM Ca^2+^ at 300 Hz, with the dashed line showing WT responses under the same conditions. Data are shown as the means±SEMs. Statistical significance was determined by t test. ns, not significant.

Given the comparable RRP size between two genotypes, we next investigated the effects of I_Ca_ with or without CDF on quantal output. EPSC trains evoked by afferent-fiber stimulation at different frequencies (100 and 300 Hz, 100ms) were recorded in WT and IM-AA KI mice. As shown in Figure 4 F and G, the magnitude of STD decreased in the KI mice at both frequencies with less steady-state depression of 24.1 ± 2.5 % versus 12.4 ± 1.5% in WT at 300 Hz (n=18 and n=17, respectively, t-test, ***p=0.0004, Figure 4H-I), and 50.6 ± 3.6 % versus 43.6 ± 3.8 % in WT at 100 Hz (n=19 and n=19, respectively, K-S test, ns, p=0.4602, t-test, Figure 4H-I). These differences can be attributed to attenuated quantal output in the absence of CDF during train stimulation in IM-AA synapses.

As the strong synaptic depression was observed while the presynaptic Ca^2+^ current facilitated at 2 mM extracellular Ca^2+^, we reasoned that the effects of IM-AA mutant on synaptic facilitation might be masked by synaptic depression. We then recorded the synaptic plasticity at a lower [Ca^2+^]_o_ of 1 mM. Indeed, the synaptic depression phenotype of WT synapse was immediately converted into synaptic facilitation at 1 mM [Ca^2+^] _o_ with maximal facilitation 197 ± 14% (n=10) at 300 Hz, and 132 ± 18.9 % at 100 Hz (n=8) trains (Figure 4J, L-M). In contrast, IM-AA synapses showed less facilitation at two frequencies with a slower time course to reach the maximal (220 ± 17 % at 300 Hz; 170 ± 15% at 100 Hz, n=8), respectively (Figure 4K, L-M). These data suggest that STF is driven by CDF-dependent and CDF-independent mechanisms, and that CDF plays an indispensable role in expanding dynamic gain range and time course of quantal output, depending on [Ca^2+^] _o_ and the intensity of repetitive activity.

### Increase in I_Ca_ by broadening presynaptic AP width normalizes initial quantal output but not short-term plasticity

Given the differences in amplitude and time course of eEPSCs between WT and IM-AA P/Q channels, we next explored if deficits in STF and STD in IM-AA synapses can be rescued by increasing the magnitude of presynaptic Ca^2+^ transient. To this end, we made use of a low dosage of tetraethylammonium (TEA) to block presynaptic voltage-gated K^+^ channels with the rationale that broadening presynaptic APs should be able to increase I_Ca_ amplitude and prolong presynaptic Ca^2+^ transient, thereby rescuing the eEPSC amplitude. To access the effect of TEA on synaptic facilitation in IM-AA KI mice, we measured the eEPSCs during a 300 Hz train before (Figure 4N) and after (Figure 4O) washing in 50 μM TEA at 1 mM and 2 mM external Ca^2+^. The amplitude of the first eEPSCs in IM-AA KI synapses were rescued to the level of WT synapses after washing in 50 μM TEA, but TEA resulted in more robust STD in both Ca^2+^ concentrations (Figure 4P,Q). It was most evident that STF in WT synapses at 1 mM Ca^2+^ failed to be restored by TEA in IM-AA KI calyces (Figure 4Q), suggesting activity-dependent synaptic facilitation depends on CDF, which is absent in IM-AA KI synapses. This data supports the conclusion that CDF embedded in IQ motif of P/Q type Ca^2+^ channels is required for determining both magnitude and polarity of short-term synaptic plasticity during sustained neuronal activities.

## Discussion

Synapse displays great variations in strength and plasticity during repetitive stimulation. The mechanisms underlying such diversity are not fully understood. In this study, we showed that the CDF of P/Q type VGCCs is crucial for the size and trajectory of frequency-dependent synaptic plasticity at calyx of Held synapse. Using a mouse model with point mutations at IQ-like motif of the P/Q-type VGCCs, we showed that this motif not only imparts CDF but also regulates its channel density at the nerve terminal. Loss of the IQ-like motif completely precluded the action of CaM superfamily proteins to P/Q-type channels ^2^ and abolished CDF of I_Ca_ in IM-AA synapse, leading to profound changes in STF and STD. Taken together, we provide direct evidence that CDF of P/Q type VGCCs plays an indispensable role in dictating the magnitude, polarity, and temporal profile of short-term synaptic plasticity, ultimately expanding the coding capacity of information transfer in neuronal networks.

### CDF and CDI in native synapses

Several studies have assessed the functional consequences on the presynaptic Ca^2+^ channel properties upon IM-AA mutations. However, limited by the direct access to presynaptic terminals, the conclusions remain ambiguous. Using hippocampal microculture, Nanou et al^6^ showed that neither the density nor the kinetics of I_Ca_ in the soma was altered upon IM-AA mutations. On the other hand, in dissociated Purkinje neurons from IM-AA cerebellum, I_Ca_ was reduced while CDF is absent under physiological conditions^21^. The discrepancy might arise from distinct VGCC subtypes in different neurons and/or their location, some of which might be dominant in somata over those in presynaptic terminals of the neurons^29^. Compartment-specific sorting mechanism must be at play in highly polarized cells like neurons to differentiate targeting of VGCC subtypes both temporally and spatially. This has been shown at the calyx of Held-MNTB synapse, whereas somatic Ca^2+^ currents are largely contributed by L-, N- and R-types of VGCCs instead of P/Q type VGCCs^30^. This suggests that characteristics of somatic Ca^2+^ channels are not sufficient to infer their behavior in presynaptic terminals. That could provide a possible explanation that only minimum Ca^2+^ dependent facilitation was detected^21^, as L-, N- and R-types Ca^2+^ channel facilitation is less prominent compared to P/Q type Ca^2+^ channel^30^. In this study, we recorded Ca^2+^ current directly from the mature calyx of Held terminals, where P/Q type Ca^2+^ channels contribute to 99% of the Ca^2+^ current^22,23^. Using AP-like depolarization steps, we observed an average Ca^2+^ current facilitation up to 25% (at 300 Hz). This degree of facilitation agrees well with the Ca^2+^ current facilitation observed previously in response to two identical AP waveform voltage-clamp commands in rat calyces of Held ^11,14^. This facilitation is ablated in IM-AA synapses, indicating the necessity and specificity of the IQ like motif in controlling CDF, rationalizing IM-AA mice as a perfect model to unravel the effect of CDF in synaptic plasticity without confounding effects of CDI.

While both CDF and CDI of P/Q type Ca^2+^ channel exist at calyx of Held synapse ^12–14^, it is possible that inactivation of I_Ca_ contributes to the synaptic plasticity during prolonged activity. However, we did not observe apparent activity-dependent inactivation of Ca^2+^ current or CDI with 200 ms train stimulation at 300 Hz using P16-21 mice (Figure 3A, C). This is supported by previous study where they investigated the contribution of Ca^2+^/CaM-dependent inactivation of I_Ca_ in synaptic depression using specific CaM inhibitor (MLCK peptide)^31^ and found the dependence of Ca^2+^/CaM inactivation of I_Ca_ declines during synapse maturation. Previous studies showing CDI have also used much longer depolarization steps (1 ms duration, from -80 mV to 0 mV) than that we used in this study (0.3 ms duration, from -80 mV to +40 mV). These two paradigms can give rise to very different profiles of local Ca^2+^ transients and global accumulation in the nerve terminals because of differences in driving force, open probability, and number of activated VGCCs. We suggest that CDF by our paradigm, which closely mimics native AP trains, plays a predominant role in regulating synaptic plasticity in the mature calyx of Held synapse.

### The role of IQ-like motif in Ca^2+^ channel sorting and gating properties

We found the density and nanoclusters of P/Q-type VGCCs are significantly reduced in the presynaptic terminals upon IM-AA mutations (Figure 1), suggesting that the interaction between CaM (and/or other CaM superfamily members) and channel protein is crucial for these channels to be targeted to the presynaptic terminal with proper abundance. Indeed, CaM has been proposed to be a constitutive subunit for Ca^2+^ channels owing to the fact that it is associated with every type of high-voltage activated Ca^2+^ channel^32^. The association in the early stage of Ca^2+^ channel biosynthesis affords the opportunity for Ca^2+^-dependent regulation of membrane targeting^33^. Deletion of preIQ3 region-a second sequence important for Ca^2+^/CaM binding in combination of IQ motif, is sufficient to completely abolish surface expression of Cav1.2 in the presence of β2a subunit ^34^. On the other hand, downstream of IQ-motif, there are multiple sites at the C-terminus of Cav2.1 proposed to interact with different action zone scaffold proteins such as RIM (Rab3-interacting molecule)^35,36^, RBP (RIM-binding proteins)^37^, and CASK (cytoskeletal matrix of the AZ)^38^, which are thought to be important to bring VGCCs to the proximity of the release sites and couple them to the release machinery. While deletions of these interaction sites on the C-terminus of the Cav2.1 surprisingly have little effect on VGCC density at presynaptic terminal^39^, suggesting other motifs are responsible for the channel targeting. Our findings imply the IQ motif as a novel interaction site in controlling the abundance of VGCC at presynaptic terminal.

We found P/Q channels with IM-AA mutations display little activity-dependent change in their gating kinetics compared to the WT, suggesting the basal activity of these channels has also changed upon loss of IQ-like motif. We speculate the change of gating properties might be due to loss of pre-association with calcium-free Calmodulin (apoCaM). It was reported that apoCaM can pre-associate with Cav2.1 and prime the channel for “Camodulation” ^32^. Interaction between L-type Ca^2+^ channels to apoCaM can enhance the baseline open probability of channel by nearly 7 fold ^40^ and the ambient CaM levels and Ca^2+^ entry through the channel could then retune channel affinity for apoCaM to further control the strength of Ca^2+^ feedback ^41^. Lower abundance of VGCCs in IM-AA mice might also change the homeostasis of the Ca^2+^ signal and consequently the set-point of presynaptic fusion machinery during development^42^.

### IQ motif-mediated CDF of Ca^2+^ channel in native synapses

CaM/IQ motif-mediated CDF has been studied over the years, and several mechanisms have been put forward. However, little is known about how CaM/IQ motif interaction modulates the Cav2.1 channel level and activity in the native synapse. As CaM, a Ca^2+^ sensor, has the capability to bind to Ca^2+^ at the range of 10^-12^M-10^-6^M^43^, its intrinsic difference in Ca^2+^ binding affinity at its N-lobe, C-lobe, and linker region gives it the flexibility in structure to dynamically regulate its target proteins in a wide range of intracellular [Ca^2+^]. This explains why CDF only reaches its plateau 20 ms after the initial stimulation at 300 Hz (Figure 3D). Perhaps it is due to the time required for an initial buildup of Ca^2+^ in the terminal to activate CaM. The timing of maximal STF matches the plateau of CDF well (Figure 4M). The delay of increasing in I_Ca_ charge perhaps also reflects a progressive augmentation of “facilitated unitary Cav2.1 channel” ^10^. It is still debated whether the facilitation of Cav2.1 is through an enhanced steady-state opening ^10^ or an accelerated activation of the Cav2.1^11^. One could predict if an accelerated activation is the case, the boost in open probability at activation should only increase in I_Ca_ amplitude without changing the waveform of the I_Ca_. Conversely, if the enhanced steady-state opening hypothesis is true, we should also observe a longer-lasting I_Ca_ decay time constant. Indeed, we found the deactivation time constant of the I_Ca_ was gradually prolonged during 300 Hz pseudo-AP stimulation, with a 7% increase in the half-width of I_Ca_ compared to the first I_Ca_ time constant in WT (Figure 3H). In contrast, this phenomenon is completely absent in IQ motif mutation (Figure 3I). Although these two mechanisms are not mutually exclusive, our results support the idea that the CaM-mediated CDF works through a prolonged deactivation model.

### Mechanism underlying CDF of Ca^2+^ channel in regulating synaptic depression and facilitation

Our results show that both STF and STD were altered upon mutations at the IQ-like motif at calyx of Held synapse, demonstrating that IM-AA mutations modify the Ca^2+^-dependent encoding of information in response to trains of AP, resulting in an altered timing and pattern of short-term plasticity. This is consistent with previous work in the neuromuscular junction^18^, hippocampal culture neuron and SC-CA1 synapse^6^, where different degrees of attenuation in STF and STD were observed upon mutating the IQ-like motif; However, these observations contradict to the findings using hippocampal CA3-CA1, PF-PC and PC-DCN synapse in IM-AA mice^21^, where STF remains unaltered and partly even enhanced. The authors reasoned the enhanced STF might be due to a reduced Ca^2+^ influx in the IM-AA KI case, which led to a lower initial Pr, thereby amplifying the tendency of facilitation. Alternatively, the contribution of CDF-dependent component to STF/STD is largely masked by that of CDF-independent component when P/Q-type channels are not dominant in these synapses using multiple types of calcium channels. At the calyx of Held synapse where P/Q-type channel exclusively mediates transmitter release, we found that Ca^2+^ influx is reduced by 35% in IM-AA calyx (Figure. 1C), so are the absolute magnitude of STF/STD. Experimental rescue of the Ca^2+^ influx using prolonged APs largely normalizes the initial quantal output from IM-AA synapses, but STF and STD were not fully rescued (Figure 4P-Q), suggesting the pivotal role of CDF of Ca^2+^ channel in expanding the dynamic range of short-term plasticity during intense synaptic activity.

## Acknowledgements

We would like to dedicate this work to the memory of Prof. William A Catterall, who personally inspired us to study calcium-dependent facilitation at the calyx of Held synapse in IMAA-KI mice generated from his lab. This work was supported by by grants from the Natural Sciences and Engineering Research Council of Canada (NSERC RGPIN-2022-05425), Canadian Institutes of Health Research (CIHR PJT-156034; PJT-505732; PJT-518084) and Tier 1 Canada Research Chair (CRC-95-232466) to L.Y.W. and The Deutsche Forschungsgemeinschaft (DFG - German Research Foundation) postdoctoral fellowship to S.C.

## Supplementary Statistics Table

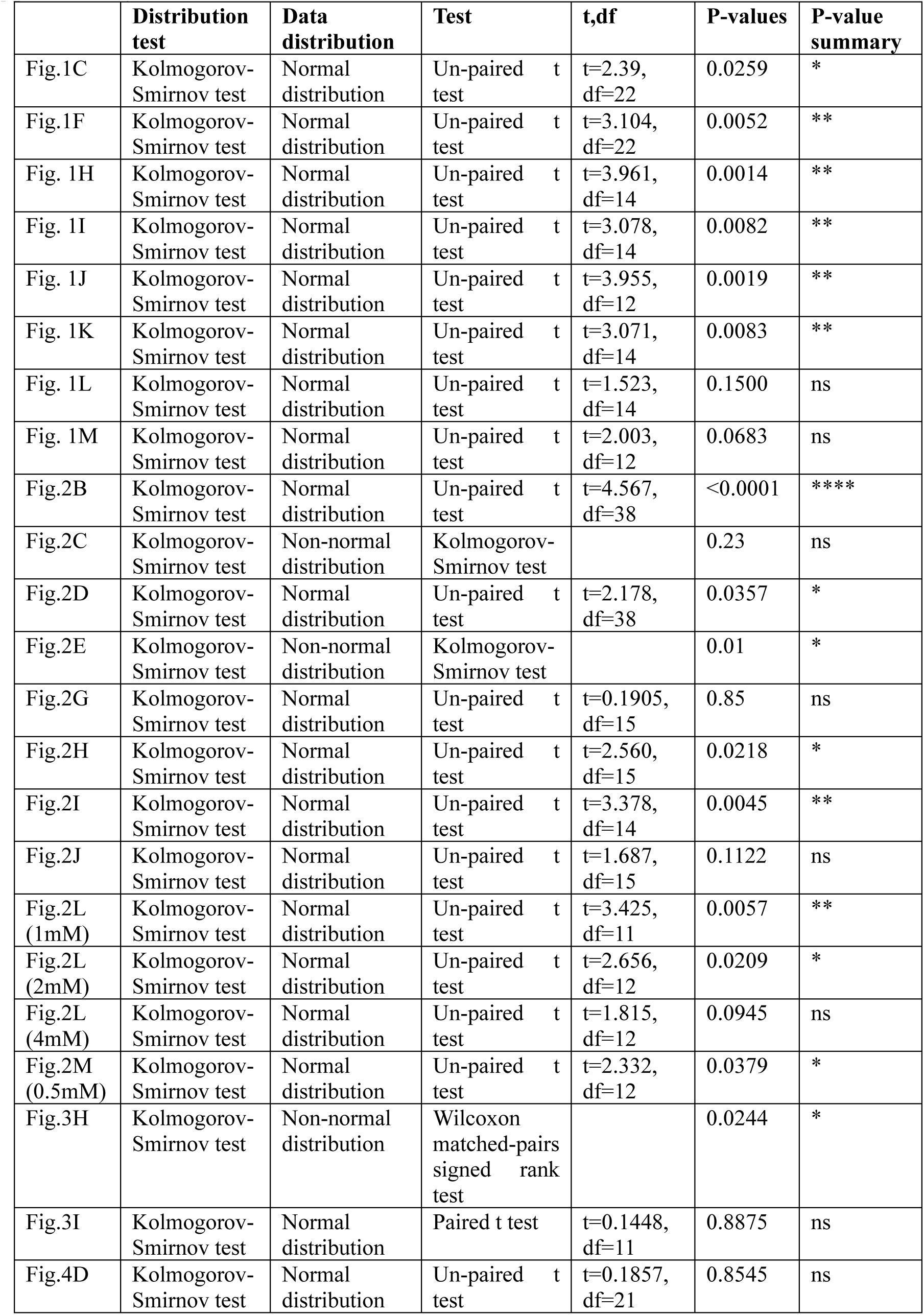
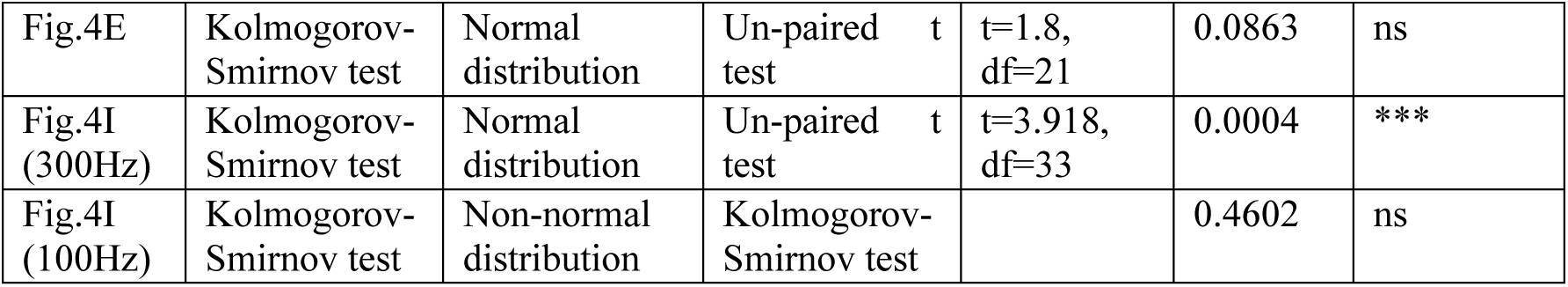

|  | **Distribution test** | **Data distribution** | **Test** | **t,df** | **P-values** | **P-value summary** |
| --- | --- | --- | --- | --- | --- | --- |
| Fig.1C | Kolmogorov-Smirnov test | Normal distribution | Un-paired t test | t=2.39, df=22 | 0.0259 | * |
| Fig.1F | Kolmogorov-Smirnov test | Normal distribution | Un-paired t test | t=3.104, df=22 | 0.0052 | ** |
| Fig. 1H | Kolmogorov-Smirnov test | Normal distribution | Un-paired t test | t=3.961, df=14 | 0.0014 | ** |
| Fig. 1I | Kolmogorov-Smirnov test | Normal distribution | Un-paired t test | t=3.078, df=14 | 0.0082 | ** |
| Fig. 1J | Kolmogorov-Smirnov test | Normal distribution | Un-paired t test | t=3.955, df=12 | 0.0019 | ** |
| Fig. 1K | Kolmogorov-Smirnov test | Normal distribution | Un-paired t test | t=3.071, df=14 | 0.0083 | ** |
| Fig. 1L | Kolmogorov-Smirnov test | Normal distribution | Un-paired t test | t=1.523, df=14 | 0.1500 | ns |
| Fig. 1M | Kolmogorov-Smirnov test | Normal distribution | Un-paired t test | t=2.003, df=12 | 0.0683 | ns |
| Fig.2B | Kolmogorov-Smirnov test | Normal distribution | Un-paired t test | t=4.567, df=38 | <0.0001 | **** |
| Fig.2C | Kolmogorov-Smirnov test | Non-normal distribution | Kolmogorov-Smirnov test |  | 0.23 | ns |
| Fig.2D | Kolmogorov-Smirnov test | Normal distribution | Un-paired t test | t=2.178, df=38 | 0.0357 | * |
| Fig.2E | Kolmogorov-Smirnov test | Non-normal distribution | Kolmogorov-Smirnov test |  | 0.01 | * |
| Fig.2G | Kolmogorov-Smirnov test | Normal distribution | Un-paired t test | t=0.1905, df=15 | 0.85 | ns |
| Fig.2H | Kolmogorov-Smirnov test | Normal distribution | Un-paired t test | t=2.560, df=15 | 0.0218 | * |
| Fig.2I | Kolmogorov-Smirnov test | Normal distribution | Un-paired t test | t=3.378, df=14 | 0.0045 | ** |
| Fig.2J | Kolmogorov-Smirnov test | Normal distribution | Un-paired t test | t=1.687, df=15 | 0.1122 | ns |
| Fig.2L (1mM) | Kolmogorov-Smirnov test | Normal distribution | Un-paired t test | t=3.425, df=11 | 0.0057 | ** |
| Fig.2L (2mM) | Kolmogorov-Smirnov test | Normal distribution | Un-paired t test | t=2.656, df=12 | 0.0209 | * |
| Fig.2L (4mM) | Kolmogorov-Smirnov test | Normal distribution | Un-paired t test | t=1.815, df=12 | 0.0945 | ns |
| Fig.2M (0.5mM) | Kolmogorov-Smirnov test | Normal distribution | Un-paired t test | t=2.332, df=12 | 0.0379 | * |
| Fig.3H | Kolmogorov-Smirnov test | Non-normal distribution | Wilcoxon matched-pairs signed rank test |  | 0.0244 | * |
| Fig.3I | Kolmogorov-Smirnov test | Normal distribution | Paired t test | t=0.1448, df=11 | 0.8875 | ns |
| Fig.4D | Kolmogorov-Smirnov test | Normal distribution | Un-paired t test | t=0.1857, df=21 | 0.8545 | ns |
| Fig.4E | Kolmogorov-Smirnov test | Normal distribution | Un-paired t test | t=1.8, df=21 | 0.0863 | ns |
| Fig.4I (300Hz) | Kolmogorov-Smirnov test | Normal distribution | Un-paired t test | t=3.918, df=33 | 0.0004 | *** |
| Fig.4I (100Hz) | Kolmogorov-Smirnov test | Non-normal distribution | Kolmogorov-Smirnov test |  | 0.4602 | ns |

### Method

#### Mice

Animal procedures have been carried out in accordance with the guidelines and protocols were approved by the Hospital for Sick Children and University of Toronto Animal Care Committees and Institutional Animal Care and Use Committee. Mice were housed in the facility certified by the Canadian Council on Animal Care. IM-AA mice in which isoleucine1913 and methionine1914 of the Cav2.1 IQ-like motif have been replaced with alanine1913 and alanine 1914 via conversion of the nucleotide sequence from ATCATG to GCCGCT, were originally generated by William Catterall’s laboratory at the University of Washington. IM-AA KI mice were backcrossed for at least eight generations into the C57BL/6N genetic background. Homozygous IM-AA KI and WT littermates in either sex at the age of P16-P21 were included in this study.

#### Slice preparation

Mice were sacrificed by decapitation. The brain was removed and placed into ice-cold solution (in mM) containing: 125 NaCl, 2.5 KCl, 10 glucose, 1.25 NaH_2_PO_4,_ 2 Na-pyruvate, 3 myo-inositol, 0.5 ascorbic acid, 26 NaHCO_3_,3 MgCl_2_, and 0.1 CaCl_2_ (pH=7.4) and sliced on a vibratome (VT1200S, Leica). Transverse brainstem slices (200-250 μm) containing the medial nucleus of trapezoid body (MNTB) were prepared. Subsequently, slices were transferred into oxygenated standard artificial cerebrospinal fluid (aCSF), 125 NaCl, 2.5 KCl, 10 glucose, 1.25 NaH_2_PO_4,_ 2 Na-pyruvate, 3 myo-inositol, 0.5 ascorbic acid, 26 NaHCO_3_,1 MgCl_2_, and 2 CaCl_2_ (pH=7.4), incubated for 1 hour at 35 °C, then stored at room temperature until use.

#### Electrophysiology

Whole-cell patch-clamp recordings were made from calyx of Held terminals and principal neurons of the NMTB using a Multiclamp 700B dual-channel amplifier (Molecular Devices, California, USA) controlled by Clampex10 software. Data were digitally sampled at 50 kHz and were filtered using a low-pass filter at 4 kHz. Principal cells embraced by the calyx of Held synapses in the MNTB were visualized by an upright microscope (Axio Examiner, Carl Zeiss) equipped with a CCD monochrome video camera (IR-1000, DAGE0MTI) or an Olympus microscope with Nomarski optics and a 60X water immersion objective. During experiments, slices were continuously perfused with normal aCSF solution. All experiments were performed at room temperature (22 °C).

#### Presynaptic recordings

Presynaptic patch pipettes were pulled from a puller (PP-830, Narishige) from borosilicate glass with filament (WPI) with a resistance of 4-5 mOhms. Patch electrodes were filled with (in mM) 110 CsCl, 0.5 EGTA, 40 HEPES, 1MgCl_2_, 2 Na-ATP, 0.5 Na-GTP, 12 phosphocreatine d(tris) salt, and 30 TEA-Cl, pH adjusted to 7.3 with CsOH (about 330 mOsm). I_Ca_ were recorded in aCSF containing 2 mM CaCl_2_, bicuculline (10 μM), strychnine (1 μM), tetridotoxin (TTX, 1 μM), TEA (10 mM) and 4-aminopyridine (4-AP, 0.3 mM) into the aCSF to block Na^+^ and K^+^ channels. The holding potential (V_h_) for presynaptic recordings was - 80 mV and the series resistance was <10 mOhms with 90 % compensation. Capacitive current and leak was subtracted online with the P/4 protocol.

#### Postsynaptic recordings

For postsynaptic recordings, the patch electrodes with 2.5-3.5 mOhms were filled with (in mM) 97.5 K-gluconate, 32.5 CsCl, 5 EGTA, 10 HEPES, 1 MgCl_2_, 30 TEA, and 3 lidocaine N-ethyl bromide (QX-314, Br-), and pH adjusted to 7.3 with KOH (about 310 mOsm). eEPSC were elicited by afferent fiber stimulation using a modified bipolar stimulator (MicroProbes for Life Science P14ST30.1B10). The afferent axons to calyx terminals were stimulated at the midline (<15V), 0.2 ms; 10-20% above threshold to avoid presynaptic AP failures) using a Master 8 stimulator (A.M.P.I) coupled to the stimulator. eEPSC were recorded in aCSF containing either 1mM or 2 mM CaCl_2_ (specified in the main text), bicuculline (10 μM), and strychnine (1 μM). To obtain Ca^2+^ dose-response curve, recordings were performed by application of aCSF containing (in mM) 0.5, 1, 1.5, 2, 4 CaCl_2_ with 2.5, 2, 1.5, 1, 0.5 MgCl_2_ respectively. The holding potential (V_h_) for postsynaptic recordings was -70 mV and the series resistance was compensated to 90%.

#### Analysis

Off-line analysis was performed using the pCLAMP10 software package (Molecular Devices), Axograph X (Axograph Scientific, Sydney, Australia), Clampfit (Molecular Devices). Individual eEPSC trains were baseline subtracted and averaged. We quantified the EPSC amplitudes recorded during the stimulus trains at 100Hz -300 Hz (100-200 ms). The size of the readily releasable pool (RRP) of synaptic vesicles was estimated from the depleting stimulus train (300 Hz, 200 ms) by back-extrapolating the time 0 ms from last 33 ms of the steady-state portion of the cumulative EPSC amplitude curves.

#### Mouse brain tissue preparation for expansion microscopy

We anesthetized P18 IM-AA KI and WT littermate mice (C57BL/6 background) with a ketamine (90mg/Kg) and xylazine (10mg/Kg) solution and perfused intracardially with 15-20 ml ice-cold, modified artificial cerebrospinal fluid (aCSF) solution containing (in mM): 2 CaCl_2_, 1 MgCl_2_, 125 NaCl, 2.5 KCl, 10 glucose, 26 NaHCO_3_, 3 myo-inositol, 1.25 NaH_2_PO_4_, 2 Na pyruvate, 0.5 ascorbic acid (pH = 7.4), 0.2mg/ml heparin, oxygenated with 95% O_2_ and 5% CO_2_. Next, 15 ml ice-cold fixative solution containing 20% acrylamide (AAm) and 4% paraformaldehyde (PFA) in 1x phosphate-buffered saline (PBS) was perfused. Brains were dissected and incubated in the fresh fixative for at least 48 hours, washed 3 times in 1x PBS at room temperature, then placed in a hypertonic 30% sucrose in PBS solution for at least 72 hours at 4°C. Brains were frozen in optimal cutting medium at -80°C, then 50 µm brainstem MNTB slices were cryosectioned (CryoStar NX70 cryostat, Thermo-Fisher), and washed 3 times with 1x PBS for 10 mins at room temperature. Tissues were stored in 1x PBS at 4°C until expansion microscopy (ExM).

#### ExM reagents

40% ultra pure acrylamide (AAm) solution was purchased from BioShop Canada (ACR006). 16% paraformaldehyde (PFA) was purchased from Electron Microscopy Sciences (15710). N,N′-(1,2-dihydroxyethylene)bisacrylamide (DHEBA) was purchased from Toronto Research Chemicals (D493225). Sodium acrylate (SA) was purchased from Sigma-Aldrich (408220). 38% (w/v) SA solutions were made in water and checked for purity according to solution color. Only solutions that were light yellow were used while bright yellow or cloudy solutions were discarded. Ammonium persulfate (APS) was purchased from Sigma-Aldrich (A3678). N,N,N′,N′-tetramethylethylenediamine was purchased from Sigma-Aldrich (T7024). Next, 10x phosphate buffered saline (10x PBS) was purchased from Thermo Fisher (70011044). Sodium dodecyl sulfate (SDS) was purchased from Sigma-Aldrich (436143).

#### Expansion microscopy

The ∼5x expansion microscopy (ExM) protocol was adapted and modified from M’Saad et al., 2025. We first incubated the MNTB slices in “inactivated” expansion gel solution (19% (w/v) SA + 10% AAm (w/v) + 0.1% (w/v) DHEBA in 1× PBS) for 45 min at 4°C on a rocker, and then incubated in “activated” expansion gel solution (19% (w/v) SA + 10% AAm (w/v) + 0.1% (w/v) DHEBA + 0.075% (v/v) TEMED + 0.075% (w/v) APS in 1× PBS) for 30 minutes at 4°C on a rocker before placing in a gelation chamber. The gelation chambers with the tissue sections were stored at room temperature for 15 minutes until moving them to a humidified chamber for 2 hours at 37 °C. The tissue-gel hybrids were carefully removed from the gelation chamber and incubated in ∼2 ml denaturation buffer (200 mM SDS + 200 mM NaCl + 50 mM Tris in MilliQ water, pH 6.8) in 15 ml Falcon tubes for 1-2 hours at room temperature. After replacing with fresh ∼2 mL denaturation buffer, the gels were incubated in a humidified chamber at 75 °C for 4 hours. The gels were then washed 3 times with 1x PBS for 30 minutes each and optionally stored in 1x PBS at 4 °C until immunohistochemistry.

#### Post-ExM immunohistochemistry

Hydrogel-embedded, 50 µm MNTB slices from P18 wildtype mice were blocked in 0.3% triton X in PBS and 10% normal goat serum for 2 hours at 4℃. The tissues were then incubated in primary antibody (polyclonal guinea pig anti-Ca2+ channel P/Q-type alpha-1A, Synaptic Systems, Cat #: 152 205 and monoclonal rabbit anti-GluA4, Cell Signaling, Cat #: 8070S; all 1:250 dilution) and antibody buffer (0.3% triton X in PBS and 10% normal goat serum) overnight at 4℃ on a rocker. Tissues were washed in 0.1% Triton X in 1x PBS 3 times for 5 minutes, 2 times for 10 minutes, and 3 times for 30 minutes at 4℃. Tissues were then incubated in secondary antibody (goat anti-ATTO647N, Sigma-Aldrich, Cat # 40839; goat anti-guinea pig Alexa Fluor 546, Invitrogen, Cat #: A-11074; all 1:250 dilution) and antibody buffer (0.3% triton X in PBS and 10% normal goat serum) overnight at 4℃ with gentle agitation. Tissues were washed in 1x PBS 3 times for 5 minutes, 2 times for 10 minutes, and 3 times for 30 minutes. Finally, to provide ultrastructural context, gels were incubated with 30 μg/mL Alexa Fluor 405 NHS ester (Invitrogen, A30000) in 100 mM sodium bicarbonate buffer for 2 hours at room temperature. Gels were then washed in 100 mM sodium bicarbonate buffer 4 times for 5 minutes, and then in 1x PBS 2 times for 10 minutes. Stained, hydrogel-embedded tissues were then stored in 1x PBS at 4℃ until imaging.

#### Confocal imaging of expanded MNTB hydrogel-tissue hybrids

Stained, hydrogel-embedded tissues were placed in 6-well plates filled with MilliQ water for the ∼5x expansion. Water was exchanged 3 times every 15 minutes, then the gels were left to expand in water overnight before the imaging session. The gels were trimmed to contain only the MNTB region for easier handling. Gels expanded ∼4.5-5× according to sodium acrylate purity. Imaged were acquired using an inverted Leica Stellaris 5 Confocal imaging system, equipped with a Leica DMI8 microscope and super-sensitive hybrid detectors (HyD). Images were acquired using a 40x/1.1 NA objective (motCORR) and samples were sequentially scanned with 404 nm, 494 nm, 552 nm, and 641 nm laser beams generated by a White Light Laser. The gel was immobilized for imaging by gently placing it on a 35 mm glass bottom dish and removing all excess water with a tissue. The camera pixel size in the x-y direction was 167 nm x 167 nm, and the step size of the z-stack was 250 nm. All acquired z-stack images contained 1 single calyx.

#### Cav2.1 and GluA4 cluster analyses

Fiji ImageJ (NIH) software was used to render 3D reconstructions of the calyx image and to adjust contrast and brightness. Each image channel was processed with a Mean convolution filter (radius = 1 pixel) to reduce noise. To quantify Cav2.1 and GluA4 clusters, we used the 3D object counter plugin in Fiji ImageJ (https://imagej.net/3D_Objects_Counter, online manual available, NIH). The Cav2.1 and GluA4 fluorescence throughout the calyx was thresholded above the background level, then each identified cluster was assigned a number, and its integrated density was measured (Fig. 1 I, L). We quantified the contact area using the region of interest manager tool (ROI, ImageJ). We measured the perimeter of the calyx that physically contacted each cluster and multiplied it by the step-size (0.250 µm). The total number of nanoclusters (Fig. 1H, K) was normalized by dividing the total by the sum of contact areas for each calyx. Overall protein fluorescence intensity of Cav2.1 and GluA4 was measured as integrated density of a maximum projection of calyx z-stack (Fig. 1J, M). Averages were expressed as mean ± S.E.M. Statistical significance was established using unpaired t tests as noted, with p < 0.05 indicating a significant difference.

#### Statistics

Data summation and statistical analyses were performed using Prism (GraphPad, California, USA). All data were first subjected to a normality test. Two-tailed paired or unpaired Student’s t tests for normally distributed data and the Kolmogorov-Smirnov test or Wilcoxon matched-pairs rank test for non-normally distributed data were then carried out. Differences between data sets were considered insignificant at P values < 0.05.

